# The PAGE Study: How Genetic Diversity Improves Our Understanding of the Architecture of Complex Traits

**DOI:** 10.1101/188094

**Authors:** Genevieve L Wojcik, Mariaelisa Graff, Katherine K Nishimura, Ran Tao, Jeffrey Haessler, Christopher R Gignoux, Heather M Highland, Yesha M Patel, Elena P Sorokin, Christy L Avery, Gillian M Belbin, Stephanie A Bien, Iona Cheng, Sinead Cullina, Chani J Hodonsky, Yao Hu, Laura M Huckins, Janina Jeff, Anne E Justice, Jonathan M Kocarnik, Unhee Lim, Bridget M Lin, Yingchang Lu, Sarah C Nelson, Sung-Shim L Park, Hannah Poisner, Michael H Preuss, Melissa A Richard, Claudia Schurmann, Veronica W Setiawan, Alexandra Sockell, Karan Vahi, Abhishek Vishnu, Marie Verbanck, Ryan Walker, Kristin L Young, Niha Zubair, Victor Acuna-Alonso, Jose Luis Ambite, Kathleen C Barnes, Eric Boerwinkle, Erwin Bottinger, Carlos D Bustamante, Christian Caberto, Samuel Canizales-Quinteroes, Matthew P Conomos, Ewa Deelman, Ron Do, Kimberly Doheny, Lindsay Fernandez-Rhodes, Myriam Fornage, Gerardo Heiss, Brenna Henn, Lucia A Hindorff, Rebecca D Jackson, Benyam Hailu, Cecelia A Laurie, Cathy C Laurie, Yuqing Li, Dan-Yu Lin, Andres Moreno-Estrada, Girish Nadkarni, Paul Norman, Loreall C Pooler, Alexander P Reiner, Jane Romm, Chiara Sabati, Karla Sandoval, Xin Sheng, Eli A Stahl, Daniel O Stram, Timothy A Thornton, Christina L Wassel, Lynne R Wilkens, Cheryl A Winkler, Sachi Yoneyama, Steven Buyske, Chris Haiman, Charles Kooperberg, Loic Le Marchand, Ruth JF Loos, Tara C Matise, Kari E North, Ulrike Peters, Eimear E Kenny, Christopher S Carlson

**Affiliations:** Stanford University, Stanford CA; University of North Carolina at Chapel Hill, Chapel Hill NC; Fred Hutchinson Cancer Research Center, Seattle WA; Vanderbilt University Medical Center, Nashville TN; Keck School of Medicine, University of Southern California, Los Angeles CA; Icahn School of Medicine at Mount Sinai, New York NY; Cancer Prevention Institute of California, Fremont CA; University of Hawaii Cancer Center, Honolulu HI; University of Washington, Seattle WA; Brown Foundation Institute for Molecular Medicine, The University of Texas Health Science Center, Houston TX; Information Sciences Institute, University of Southern California, Marina del Rey CA; Human Genetics Center, School of Public Health, The University of Texas Health Science Center, Houston TX; University of Hawaii, Honolulu HI; Johns Hopkins University, Baltimore MD; NIH National Human Genome Research Institute, Bethesda MD; Ohio State Medical Center, Columbus OH; NIH National Institute on Minority Health and Health Disparities, Bethesda MD; University of Vermont College of Medicine, Burlington VT; Rutgers University, New Brunswick NJ; Escuela Nacional de Antropologia e Historia, Mexico; University of Colorado, Anschutz Medical Campus, Denver CO; Instituto Nacional de Medicina Genómica, Mexico; The Pennsylvania State University, University Park, PA; University of California, Davis CA; National Laboratory of Genomics for Biodiversity (UGA-LANGEBIO), CINVESTAV-Irapuato, Mexico; Frederick National Laboratory, Frederick MA

**Author notes:** Shared first authorship (Genevieve L Wojcik, Mariaelisa Graff, Katherine K Nishimura, Ran Tao, Jeffrey Haessler, Christopher R Gignoux, Heather M Highland, Yesha M Patel). Shared senior authorship (Steven Buyske, Chris Haiman, Charles Kooperberg, Loic Le Marchand, Ruth JF Loos, Tara C Matise, Kari E North, Ulrike Peters, Eimear E Kenny, Christopher S Carlson). Corresponding authorship (Eimear E Kenny, Christopher S Carlson).

## Abstract

Genome-wide association studies (GWAS) have laid the foundation for investigations into the biology of complex traits, drug development, and clinical guidelines. However, the dominance of European-ancestry populations in GWAS creates a biased view of the role of human variation in disease, and hinders the equitable translation of genetic associations into clinical and public health applications. The Population Architecture using Genomics and Epidemiology (PAGE) study conducted a GWAS of 26 clinical and behavioral phenotypes in 49,839 non-European individuals. Using strategies designed for analysis of multi-ethnic and admixed populations, we confirm 574 GWAS catalog variants across these traits, and find 38 secondary signals in known loci and 27 novel loci. Our data shows strong evidence of effect-size heterogeneity across ancestries for published GWAS associations, substantial benefits for fine-mapping using diverse cohorts, and insights into clinical implications. We strongly advocate for continued, large genome-wide efforts in diverse populations to reduce health disparities.

---

A significant bias has been noted in the field of genome-wide association studies (GWAS), with the vast majority of discovery efforts conducted in populations of European ancestry. ^1–3^ In contrast, individuals of African or Latin American ancestry accounted for only 4.2% of samples analyzed. (**Extended Data Figure 1**) In light of differential genetic architecture that is known to exist between ancestral populations, there is a legitimate concern that bias in representation might exacerbate existing disease and health care disparities for several reasons. A) critical variants will be missed if they are low allele frequency or absent in European-descent groups, and B) effect sizes and risk prediction scores derived from one population may not accurately extrapolate to other populations. Indeed, recent seminal papers have demonstrated that some genetic predictors are restricted to certain ancestries, and thus may partially explain risk differences among populations. ^4–10^ Additionally, as the field shifts its attention towards low frequency variants, which are more likely to be population-specific, we can no longer rely on the transferability of findings from one population to another, a complication that has also been observed with some common variants. ^11,12^ In the United States where minority populations have a disproportionately higher burden of chronic conditions ^13^, the lack of representation of diverse populations in genetic research will result in inequitable access to precision medicine for those with the highest burden of disease.

Many factors contribute to the Euro-centric bias in genetic research ^3^, including a paucity of studies recruiting diverse populations, and a lack of statistical tools specifically developed to leverage multi-ethnic and admixed study populations. ^14^ However, recent advancements in statistical analyses and genotyping technologies have addressed many methodological concerns, removing barriers that had previously made researchers reluctant to recruit and analyze heterogeneous samples. Likewise, over the past decade the Population Architecture using Genomics and Epidemiology (PAGE) study has catalyzed the genomics community by building the next generation of arrays, methods, resources, and guidance for the interrogation of the genetics of underrepresented populations. ^15,16^ Here we report on our study of 49,839 individuals of non-European ancestry (see **Supplementary Info Section 1** for demographic details) and describe results from our multi-ethnic GWAS analysis across 26 traits and diseases. Using a purpose-designed array and statistical methodology tailored for diverse populations, our study allowed us to systematically test for population differences in effect size for risk variants reported in the current GWAS literature, as well as to screen for potentially novel alleles within previously reported loci (i.e. secondary signals). We further explore trends between our results using a genetically diverse sample, relative to meta-analyses of an equivalent number of European individuals sampled from the UK Biobank, showing strong evidence of effect-size heterogeneity across ancestries for published GWAS associations, and substantial benefits for fine-mapping using diverse cohorts. Lastly, we report on the global patterns of allele frequencies of clinically relevant variants and illustrate the many advantages of genomics research in ancestrally diverse populations.

## Results

### Unique Methodological Challenges are Inherent to Multi-ethnic Studies

PAGE was developed by the National Human Genome Research Institute and the National Institute on Minority Health and Health Disparities to conduct genetic epidemiological research in ancestrally diverse populations within the United States, drawn from three major population-based cohorts (Hispanic Community Health Study/Study of Latinos (HCHS/SOL), Women’s Health Initiative (WHI), and Multiethnic Cohort (MEC)) and one metropolitan biobank (Bio*Me*). Genotyped individuals self-identified as Hispanic/Latino (N=22,216), African American (N=17,299), Asian (N=4,680), Native Hawaiian (N=3,940), Native American (N=652), or Other (N=1,052, primarily South Asian or mixed heritage, as well as participants who did not identify with any of the available options; **Supplementary Table 1, Supplementary Info Section 1**). Using detailed phenotype data collected and harmonized across studies (**Supplementary Info Section 2**), we present genetic association results for 26 phenotypes related to inflammation, diabetes, hypertension, kidney function, cardiac electrophysiology, dyslipidemias, anthropometry, and behavior/lifestyle (smoking and coffee consumption). Our study explicitly solves multiple challenges inherent to multi-ethnic studies, including: (1) an initial study design with prioritization of diverse participants across pre-existing studies, (2) the development of a novel genotyping platform to capture global variation with the Multi-Ethnic Genotyping Array (MEGA) **(Supplementary Info Section 3)**, (3) analysis in a complex population structure **(Supplementary Info Section 4)**, (4) statistical modeling for association testing accounting for population structure, and (5) downstream interpretations in the context of global variation. In particular, groups have highlighted the challenges of joint analysis across multiple ethnicities, preferring meta-analysis approaches that analyze populations separately ^17,18^. We show that a joint analysis does not increase type 1 error, using tools (SUGEN, GENESIS) that explicitly model population structure, admixture, relatedness between individuals and population-specific genetic heterogeneity. ^19–23^**; Supplementary Info Section 5).** We provide specific recommendations for all of these components of a genetic study in the Supplementary Information.

In brief, 49,839 multi-ethnic individuals were genotyped on MEGA (Illumina, see **Supplementary Info Section 3**), with 1,402,653 loci passing quality control filters, and subsequently imputed to 39,723,562 variants. Sample sizes ranged from 9,066 to 49,796 individuals between the 26 traits, so variants with an effective N (effN) greater than 30 were tested for association, yielding 11.6-34.6 million unique variants per trait (details in **Supplementary Table 1**). Association models were adjusted for known phenotype-specific trait covariates, as well as the top 10 principal components of genetic ancestry, indicators for study, and self-identified race/ethnicity. (**Supplementary Info Section 6**) The primary results presented are from SUGEN (GENESIS results are provided in the **Supplementary Tables 2 and 4** for comparison). For comparison against traditional multi-ethnic approaches and to assess heterogeneity by ancestry, we also conducted analyses stratified by self-identified race/ethnicity and subsequently combined in a meta-analysis (**Supplementary Table 3).** All known variants conditioned on by trait in the conditional analyses are provided in **Supplementary Table 5**.

### New Loci Found across 26 Phenotypes, Plus Additional Signals at Previously-Discovered Loci

Since the majority of GWAS have been conducted in primarily European-descent populations, we hypothesized that the examination of underrepresented populations would reveal ancestry-specific associations that European-centric studies were unable to detect. The traditional P<5×10^−8^ was used for common variants (MAF>5%), and a more conservative threshold of P<3×10^−9^ for low frequency and rare variants ^24^. Novel findings were defined as signal reaching these thresholds after conditioning on all previously reported variants on that chromosome, more than 1 megabase (Mb) away from a known locus. We found 16 novel variants at the frequency-specific thresholds for significance, as well as 11 low-frequency loci with suggestive associations (P<5×10^−8^). (**Figure 1, Table 1, Supplementary Tables 2-3**) A deeper discussion of the biological implications of these novel findings can be found in **Supplementary Info Section 8**, along with candidate genes and candidate functional polymorphisms.

**Figure 1:**
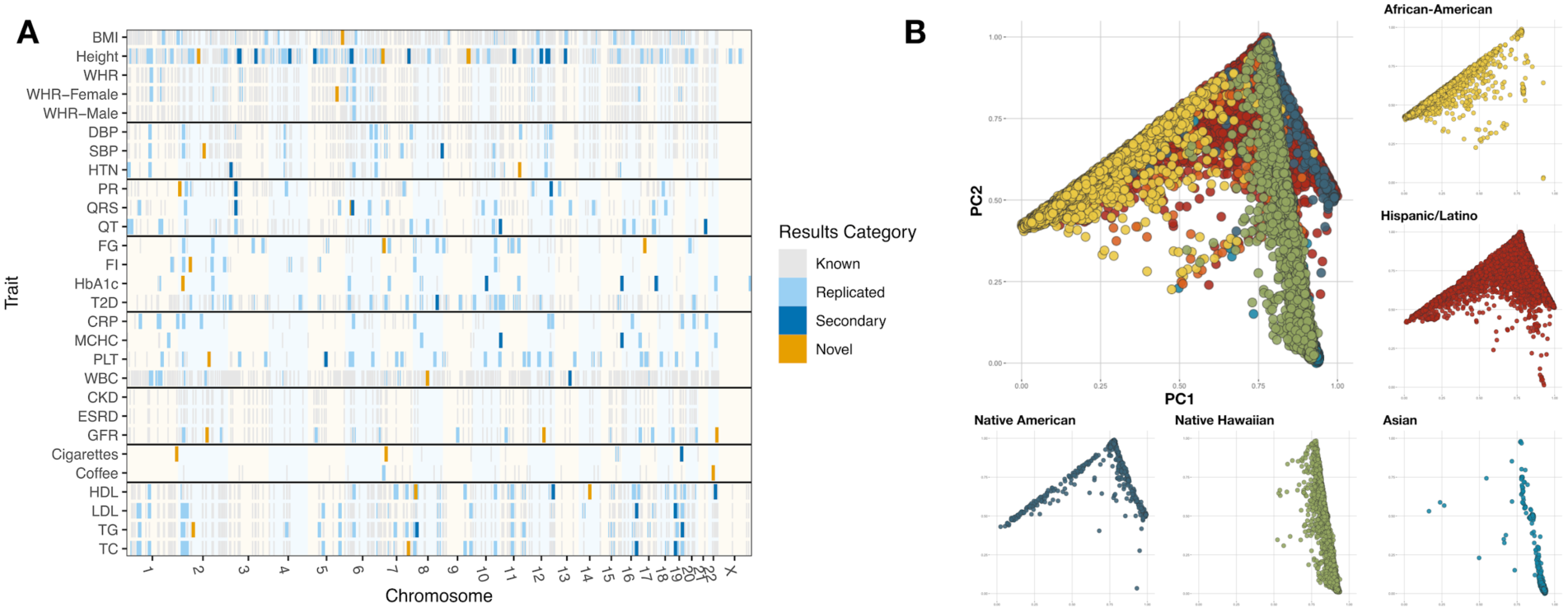
Inclusion of multi-ethnic samples enables discovery and replication in GWAS. (A) There are 8,979 previously reported trait-variant pairs, of which 1,444 replicated at a by-trait Bonferroni adjusted significance level. In addition, we found 27 novel trait-variant pairs and 38 secondary signal pairs that remained after adjusting for known variants. (B) The population substructure present in the multi-ethnic sample of PAGE revealed complex patterns preventing meaningful stratification. Here we show principal component (PC) 1 and 2 to show major patterns of variation, stratified by self-identified race/ethnicity. Individuals denoted by orange self-identified as ‘Other’. BMI = body mass index, WHR = waist-hip ratio, DBP/SBP = diastolic/systolic blood pressure, HTN = hypertension, PR/QRS/QT = PR/QRS/QT interval, HbA1c = glycated hemoglobin, FG/FI = fasting glucose/insulin, T2D = type II diabetes, CRP = C-reactive protein, MCHC = mean corpuscular hemoglobin concentration, PLT = platelet count, WBC = white blood cell count, CKD = chronic kidney disease, ESRD = end-stage renal disease, GFR = estimated glomerular filtration rate, Cigarettes = cigarettes per day, Coffee = coffee consumption, HDL/LDL = high/low-density lipoproteins, TG = triglyceride, TC = total cholesterol.

**Table 1.** GWAS Catalog heterogeneity by Trait, including number of novel and secondary findings.

|  | Largest GWAS catalog discovery population<sup>1</sup> |  |  |  |  | GWAS catalog tagSNPs |  |  | best PAGE tagSNPs |  | Novel Loci (count)<sup>6</sup> | Secondary Loci (count)<sup>6</sup> |
| --- | --- | --- | --- | --- | --- | --- | --- | --- | --- | --- | --- | --- |
| Phenotype | European | East Asian | African | Hispanic/Latino | PAGE | Unique | P<5x10<sup>-8</sup> | Het.<sup>4</sup> | P<5x10<sup>-8</sup> | Het.<sup>5</sup> |  |  |
| Inflammatory Traits |  |  |  |  |  |  |  |  |  |  |  |  |
| CRP | 66,185 | 10,112 | 8,280 | 3,548 | 28,537 | 82 | 38 | 7 | 16 | 1 | 0 | 0 |
| WBC | 19,509 | 33,231 | 16,388 | - | 28,534 | 27 | 10 | 5 | 11 | 3 | 1 | 1 |
| MCHC | 62,553 | - | 16,485 | - | 19,803 | 21 | 9 | 1 | 5 | 0 | 0 | 2 |
| Platelet Count | 48,666 | 14,806 | 7,943 | 12,491 | 29,328 | 92 | 23 | 0 | 28 | 0 | 1 | 1 |
| Lipid Traits |  |  |  |  |  |  |  |  |  |  |  |  |
| HDL | 99,900 | 12,545 | 7,917 | 4,383 | 33,063 | 244 | 71 | 8 | 21 | 1 | 2 | 2 |
| LDL | 94,595 | 12,545 | 7,861 | 4,383 | 32,221 | 192 | 46 | 12 | 18 | 0 | 0 | 2 |
| TG | 96,598 | 12,545 | 7,601 | 4,383 | 33,096 | 179 | 75 | 29 | 16 | 1 | 1 | 2 |
| TC | 100,184 | 8,344 | 6,480 | 4,383 | 33,185 | 166 | 31 | 4 | 20 | 0 | 1 | 2 |
| Lifestyle Traits |  |  |  |  |  |  |  |  |  |  |  |  |
| Cigarettes/Day Excluding Nonsmokers | 74,035 | 11,696 | 32,389 | - | 15,862 | 12 | 0 | 0 | 3 | 0 | 2 | 1 |
| Coffee Cups/Day | 91,462 | - | - | - | 35,902 | 16 | 3 | 1 | 3 | 0 | 1 | 0 |
| Glycemic Traits |  |  |  |  |  |  |  |  |  |  |  |  |
| HbA1c | 46,368 | 17,290 | - | - | 11,178 | 29 | 8 | 1 | 9 | 0 | 1 | 3 |
| Fasting Insulin | 51,750 | 7,696 | 1,040 | 229 | 21,596 | 34 | 0 | 0 | 3 | 0 | 1 | 0 |
| Fasting Glucose | 58,074 | 24,740 | 2,029 | 4,176 | 23,963 | 55 | 15 | 3 | 7 | 0 | 2 | 0 |
| Type II Diabetes<sup>2</sup> | 12,171/<br>56,862 | 15,463/<br>26,183 | 1,264/<br>5,678 | 3,848/<br>4,366 | 14,075/<br>31,752 | 286 | 28 | 2 | 13 | 0 | 0 | 1 |
| Electrocardiogram Traits |  |  |  |  |  |  |  |  |  |  |  |  |
| QT Interval | 71,061 | 6,805 | 13,105 | - | 17,348 | 183 | 39 | 1 | 11 | 0 | 0 | 2 |
| QRS Interval | 60,255 | 6,085 | 13,031 | - | 17,052 | 63 | 9 | 3 | 12 | 0 | 1 | 2 |
| PR Interval | 28,517 | 6,085 | 13,415 | - | 17,428 | 154 | 19 | 1 | 10 | 0 | 1 | 2 |
| Blood Pressure Traits |  |  |  |  |  |  |  |  |  |  |  |  |
| Systolic Blood Pressure | 74,064 | 31,516 | 29,378 | - | 35,433 | 74 | 2 | 0 | 4 | 0 | 1 | 1 |
| Diastolic Blood Pressure | 74,064 | 31,516 | 29,378 | - | 35,433 | 81 | 2 | 0 | 4 | 0 | 0 | 0 |
| Hypertension | 74,064 | 31,516 | 29,378 | - | 49,158 | 111 | 0 | 0 | 2 | 0 | 1 | 1 |
| Anthropometric Traits |  |  |  |  |  |  |  |  |  |  |  |  |
| Waist-to-hip Ratio<sup>3</sup> | 142,762 | 39,869 | 19,744 | 3,484 | 33,904 | 94 | 5 | 0 | 6 | 0 | 1 | 0 |
| Height | 253,288 | 36,227 | 20,427 | - | 49,781 | 698 | 99 | 42 | 93 | 18 | 5 | 13 |
| Body Mass Index | 236,781 | 82,438 | 39,144 | 3,484 | 49,335 | 572 | 41 | 12 | 13 | 0 | 1 | 0 |
| Kidney Traits |  |  |  |  |  |  |  |  |  |  |  |  |
| eGFR by CKD Epi Equation | 133,413 | 23,536 | 16,840 | 16,325 | 27,900 | 135 | 1 | 0 | 5 | 0 | 3 | 0 |
| Average | 90,953 | 20,953 | 14,710 | 5,570 | Total | 3356 | 548 | 194 | 333 | 24 | 27 | 38 |

<sup>1</sup> Only includes studies indexed in the GWAS Catalog on December 31, 2016
<sup>2</sup> Cases/Controls
<sup>3</sup> Includes pooled and sex-stratified studies / results
<sup>4</sup> $P < 8.71 \times 10^{-5}$ for genotype:PC interaction in PAGE, adjusting for multiple tests (0.05/574)
<sup>5</sup> $P < 1.50 \times 10^{-4}$ for genotype:PC interaction in PAGE, adjusting for multiple tests (0.05/333)
<sup>6</sup> Significant loci have $P < 5 \times 10^{-8}$ after conditioning on all known loci from the literature

We also identified novel, independent signals (secondary variants) within known loci, further enriching our understanding of the genetic architecture of traits. Of 8,979 known variant-trait combinations (involving 3,322 unique variants, some reported for multiple traits), 1,444 replicated at P<0.05 significance threshold, Bonferroni-corrected by trait. To test for secondary signals, we screened for statistical associations located within one Mb of a previously known variant-trait combination, that remained genome-wide significant after adjusting for all known variants in the “adjusted” model (**Supplementary Table 3**). As before, we applied MAF-specific P-value thresholds to the data, and identified 25 significant secondary loci at common variants (*P*_cond_<5×10^−8^), 7 significant loci at low frequency or rare variants (*P*_cond_<3×10^−9^), and 6 suggestive associations with low frequency or rare variants (*P*_cond_ between 3×10^−9^ and 5×10^−8^). If the secondary signal represents a statistically independent association, then we would expect no net change in the strength of the association between unadjusted and adjusted models, and that the known variants are in weak LD with the secondary SNPs. Roughly half of these secondary variants are consistent with an independent association (r^2^<0.2 with any known SNP). (**Supplementary Info Section 8)** These results demonstrate that at a notable fraction of known loci (14%) there is significant secondary association in PAGE, after accounting for all previously reported SNPs.

To tease apart the influence of specific ancestral components on the 27 novel and 38 secondary loci, we calculated the correlation between the risk allele genotype and each of the first ten principal components (PCs) (**Extended Data Figure 3**). These correlations reveal population structure underlying many of our novel and secondary findings, in which there are population differences in allele frequencies for the risk alleles. Most notably, the risk allele for a novel finding for cigarettes per day among smokers on chromosome 1 (rs182996728; *P*=3.1×10^−8^) was found to show significant correlation with PC4, which represents Native Hawaiian/Pacific Islander ancestry. While this variant is monomorphic or rare in most populations, it is found at 17.2% within Native Hawaiian participants, where the signal is strongest (*P_stratified_*=2.28×10^−6^). An additional example is shown with the 5 novel and secondary SNPs highly correlated with PC6, which are related to height and found to be at higher frequencies in 1000 Genomes within a subgroup of populations within East Asia, such as Japanese or Vietnamese. That our findings exhibit substantial variability in allele frequencies further illustrates a need for the inclusion of diverse populations.

### Effect Size Heterogeneity across Ancestries at Known GWAS Loci

In general, GWAS identify loci where one or more SNPs show significant association with the trait of interest. However, GWAS do not lead directly to the identification of the causal or functional variant (fSNP), which generally is in strong linkage disequilibrium (LD) with surrogate associated SNPs (tagSNPs). LD can vary between populations, so a tagSNP in perfect LD with the fSNP in one population may be in weak LD in a different population. This can lead to inconsistent estimates of the effect sizes between populations (and therefore effect size heterogeneity) if the tagSNP (instead of the causal fSNP) is used for effect size calculations. This has been explored previously in a disease-specific manner. 25–29 Because European-descent individuals are overrepresented in GWAS discovery populations and have different LD structures than other populations, we hypothesized that effect size heterogeneity between populations may exist for many previously reported tagSNP associations.

To test this hypothesis, we measured the extent of effect heterogeneity in PAGE’s multi-ethnic study population for variants previously reported to the GWAS Catalog, a compilation of findings from published GWAS. ^30^ We replicated (P<5×10^−8^) a total of 574 known tagSNPs in 261 distinct genomic regions, out of the 3,322 unique GWAS Catalog variants previously reported for our 26 traits **(Supplementary Table 4)**. However, 132 of these known tagSNPs had significant evidence of effect heterogeneity by genetic ancestry (SNPxPC *P*=8.71×10^−5^), a conservative estimate given limitations of statistical power. In 77% of these regions the strongest signal was not the previously reported GWAS Catalog variant ^30^. A complementary approach to testing for effect heterogeneity between ancestries involves directly comparing the effect sizes in PAGE against previously reported effect sizes. As suggested by Marigorta and Navarro ^31^, dividing the z-score by the square root of the number of individuals analyzed for each trait yields a dimensionless estimate of effect size (z’), and the slope of a regression on this statistic can detect systematic differences in effect size between populations. Overall, we observed a slope of 0.75 (95% CI: 0.70, 0.79), indicating that PAGE effect sizes tend to be significantly weaker than previous reports (null slope of 1). We also compared z’ from prior reports against z’ in PAGE African-Americans and Hispanics/Latinos separately (**Figure 2A)**, to assess the influence of ancestry proportions. We observed a markedly stronger reduction of effect size in African Americans (*r*=0.45; 95% CI: 0.39, 0.51) than in Hispanics/Latinos (*r*=0.81; 95% CI: 0.756, 0.86), which is most consistent with truly differential effect sizes between ancestries at the originally reported variants, rather than a winner’s curse, which would be expected to affect findings in all ethnicities equally, regardless of ancestral proportions. These results are consistent with multi-ethnic analyses differentially tagging underlying functional variation at a large portion of reported GWAS catalog loci, as opposed to truly divergent underlying fSNP effect sizes.

**Figure 2:**
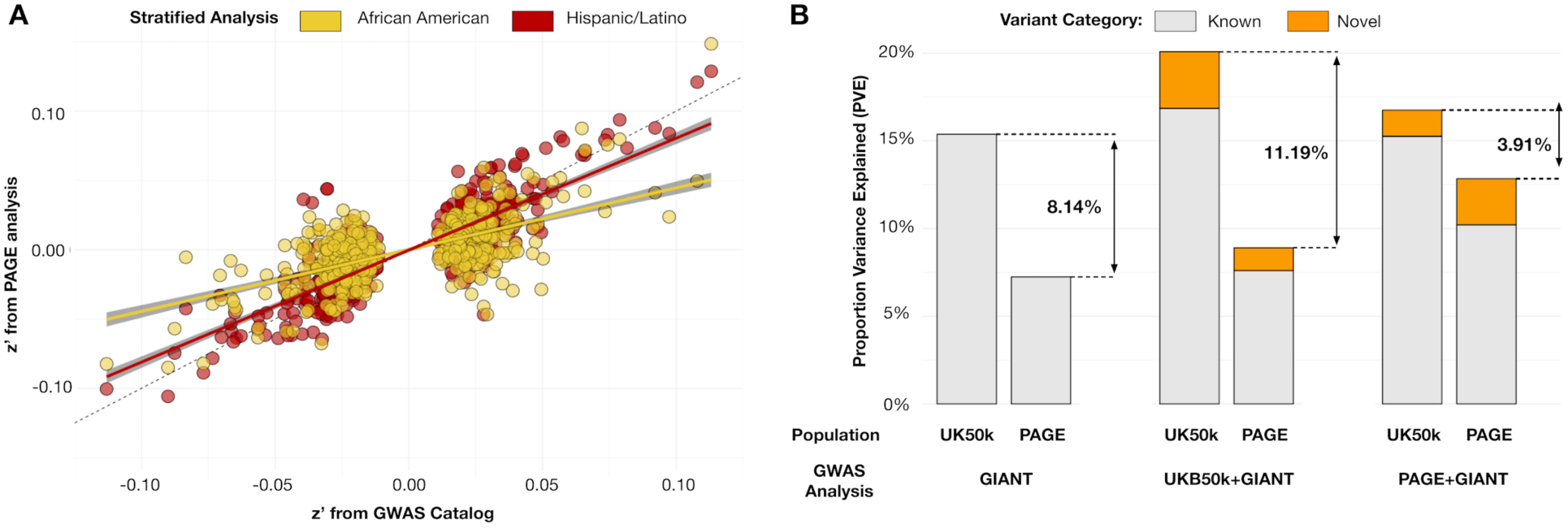
Meta-analysis with multi-ethnic samples decreases gap in proportion variance explained for height. Within each analysis, we identified the SNP with the smallest p-value in each locus, and PVE was calculated using the estimated effect size from this set of tag SNPs (left: GIANT-only GWAS, center: UKB50k+GIANT meta-analysis, bottom: PAGE+GIANT meta). PVE was estimated independently for the UKB50k (White British) and PAGE (multi-ethnic) samples, and was summed across two categories of locus: known (blue) and novel (red). The gap in PVE with previously-reported loci from GIANT (8.14%) is exacerbated with the inclusion of 50,000 more European-descent individuals to 11.19%. However, it is narrowed dramatically with the inclusion of 50,000 multi-ethnic samples to 3.91%.

### Multi-Ethnic Meta-Analysis Refines Known Loci and Reduces Disparity in Missing Heritability

Meta-analysis combining previously reported results with new GWAS data is a useful approach that simultaneously places novel findings into the context of known loci, as well as identifies novel loci that were not significant in either analysis. To quantify the value of including multi-ethnic populations in GWAS, we used published data from GIANT (a study of >250,000 individuals of European descent for anthropometric traits ^32,33^), for a meta-analysis with either PAGE (~50,000 multi-ethnic individuals) or 50,000 randomly sampled “White-British’’ individuals from the UK Biobank (UKB50k). We conducted stratified GWAS for both PAGE and UKB50k with analogous models for BMI and height, adjusting for population substructure, age, population, and sex. Both were combined in a meta-analysis with GIANT using a fixed-effect model.

Meta-analysis with GIANT resulted in many more novel findings for height (PAGE+GIANT: 82, +UKB50k+GIANT: 107) than analysis of PAGE or UKB50k alone (8 and 1 novel loci, respectively; **Extended Data Table 1**). While the number of novel loci is indicative of new insights into trait biology, understanding the proportion of variance explained (PVE) by each locus has potentially significant consequences for personalized medicine ^34^. The proportion of variance explained with loci previously reported by GIANT for height ^33^ (**Figure 2B**, top pair of bars) reveals a gross disparity, with more than twice the PVE using UKB50k summary statistics (15.4%) as in the multi-ethnic PAGE (7.2%). Meta-analysis of GIANT with 50,000 more Europeans exacerbates this disparity, with 19.2% and 8.3% PVE in UKB50k and PAGE, respectively. Even at previously known loci in GIANT, improved PVE was observed in UKB50k (1.5% gain) relative to PAGE (0.4% gain), consistent with European-specific fine-mapping. In contrast, when we use the genome-wide significant loci from the GIANT+PAGE multi-ethnic meta-analysis, the PVE gap narrows, with 16.1% and 12.0% PVE in UKB50k and PAGE, respectively. This closing of the gap comes both from the identification of novel loci with a larger PVE in PAGE, as well as from a improvement in PVE at known loci in PAGE (3.0%). Results for BMI are consistent with results for height, albeit with a substantially smaller proportion of variance explained, which is expected for a trait with lower heritability. For BMI (**Supplementary Figure 13)** the addition of 50,000 more Europeans exacerbated the gap in PVE, with nearly twice as much of the variance explained in the UKB50k (2.7%) as in PAGE (1.7%). In contrast, meta-analysis with the multi-ethnic PAGE cohort (GIANT+PAGE) improved PVE in the multi-ethnic population (2.1% in UKB50k, 2.8% in PAGE).

Meta-analysis results can also be used to fine-map associations at known loci, which can be important in identification of the functional polymorphism causing a statistical association. To assess the impact of a multi-ethnic cohort on fine-mapping precision, we compared the 95% credible sets for 390 associated regions reported by GIANT for height, as well as the 93 associated regions for BMI. We then calculated the 95% credible sets for the PAGE+GIANT and UKB50k+GIANT analyses. For height we found that on average, an additional 50,000 individuals decreased the credible sets from 11.94 SNPs in GIANT to 11.01 SNPs in UKB50k+GIANT (*P*=0.37), demonstrating limited added value **(Figure 3A).** However, the addition of 50,000 multi-ethnic individuals significantly decreased the 95% credible sets to 9.68 SNPs (*P*=0.01) Additionally, the posterior probabilities of the top ranked SNP within the credible sets was significantly higher in the PAGE+GIANT meta-analysis compared to the GIANT analysis (*P*=1.9×10^−6^) and UKB50k+GIANT analysis (*P*=3.2×10^−3^; **Figure 3B).** The addition of the UKB50k to GIANT did not significantly improve the top posterior probability (*P*=0.09). For example, the intronic variant rs11880992 in *DOT1L* was previously identified by GIANT to be associated with increased height (P=7×10^−28^) ^33^. The 95% credible sets for GIANT alone and GIANT+UKB50k contained the same four SNPs, all in high LD with each other (r^2^>0.99). (**Figure 3C-E**) However, GIANT+PAGE meta-analysis was able to narrow the 95% credible set to one SNP due to low LD between these SNPs in African-American and Hispanic/Latino populations. While trends were consistent, none of these analyses yielded significant results for BMI (*P*>0.05), likely due to the small number of regions analyzed (N=91; **Supplementary Figure 14)**

**Figure 3:**
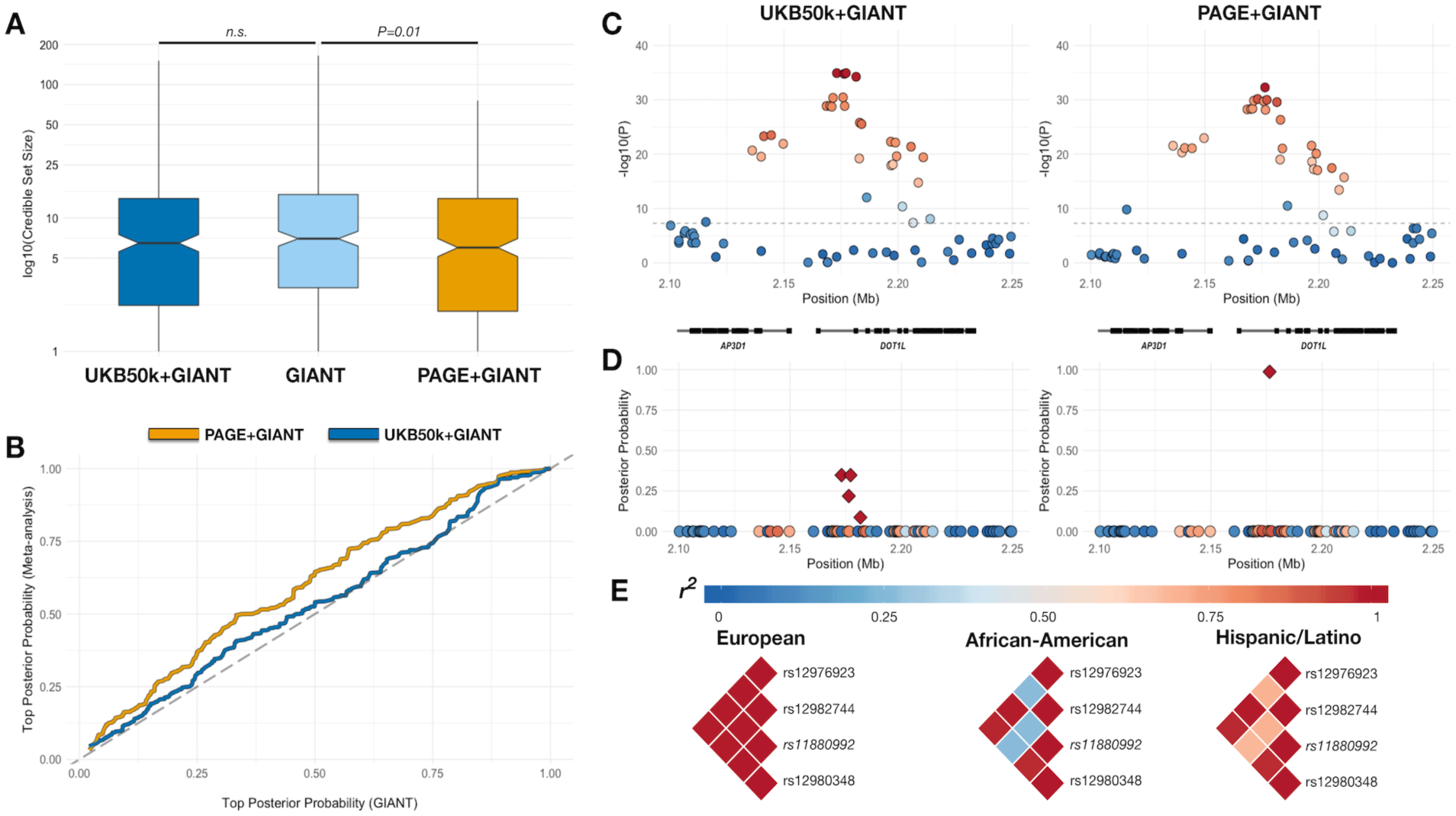
(A) Comparison of 95% credible sets for height, comparing GIANT alone to UKB50k+GIANT (P>0.05) and PAGE+GIANT (*P*=0.01). (B) Top posterior probability from each 95% credible set for height, comparing GIANT to UKB50K+GIANT and PAGE+GIANT. (C) Example of a height locus from GWAS (rs11880992), LD from weighted matrix from meta-analysis. (D) Posterior probabilities for this signal with credible set in indicated by diamond shape. (E) LD (r^2^) for the original 95% credible set from GIANT results stratified by populations.

### Relevance of Multi-ethnic Genetic Variation to Clinical Care

Concepts of diversity, genetic drift and allele frequency not only impact disease associations and predictions, but can also identify medically actionable variants. We examined the worldwide distribution of several medically actionable variants that were designed onto MEGA ^35^, and demonstrate an association between *HBB* (rs334) and HbA1c levels (P_cond_=6.87×10^−31^; N=11,178), with the majority of the signal from Hispanic/Latinos (P=7.65×10^−27^; N=10,408; MAF=0.01) and African Americans (P=5.62×10-4; N=559; MAF=0.06). The lead SNP, rs334, is a missense variant in *HBB*, which encodes the adult hemoglobin beta chain and is known for its role in sickle cell anemia. Although this association was recently reported in African Americans ^36^, this is the first time this association with HbA1c levels has been reported in Hispanic/Latinos with admixed European, African, and Native American ancestry. Hemoglobin genetic variants are also known to affect the performance of some HbA1c assays ^37–39^, potentially leading practitioners to incorrectly believe that a patient has achieved glucose control. This could leave the patient more susceptible to type II diabetes (T2D) complications. Alternative long-term measures of glucose control that are not impacted by hemoglobin variants, such as the fructosamine test, should be considered for sickle cell carriers being evaluated for T2D. This result illustrates how ancestry-specific findings may be transferable to other groups that share components of genetic ancestry; in this case the African ancestry present in both African Americans and some Hispanic/Latinos.

We also investigated the HLA-B*57:01 allotype, which interacts with the HIV drug abacavir to trigger a potentially life-threatening immune response^40–43^. The FDA recommends screening all patients for *HLA-B*57:01*, prior to starting abacavir treatment. ^44^ The rs2395029 (G) variant in *HCP5,* a near perfect tag of *HLA-B*57:01* (in Europeans), is used to screen for abacavir hypersensitivity.^45^ The tag SNP utility remains relatively high (r ~ 0.92, ^46^) across globally diverse populations in the 1000 Genomes Project. Using PAGE and Global Reference Panel samples, we show that risk allele frequencies for rs2395029 rise above 5% in multiple large South Asian populations, and rise above 1% within some, but not all, admixed populations with Native American ancestry (**Figure 4**). PAGE allele frequencies can therefore aid in expanding the reach of precision medicine to encompass individuals of diverse ancestry, particularly when combined with other studies.^47,48^. However, as previously described, if differential LD patterns are found in different populations then the utility of the rs2395029 tag SNP as a screen may not be universal in all populations. As an example, out of 152 Southern African KhoeSan individuals previously typed, 3 were identified with the *HLA-B*57:01* allotype but not the *HCP5* tag SNP, suggesting a weaker association in that population. ^49^ This argues for the need for discovery genomics programs that encompass broad and fine-scale diversity, both for common traits as described here, as well as for clinical variants.

**Figure 4:**
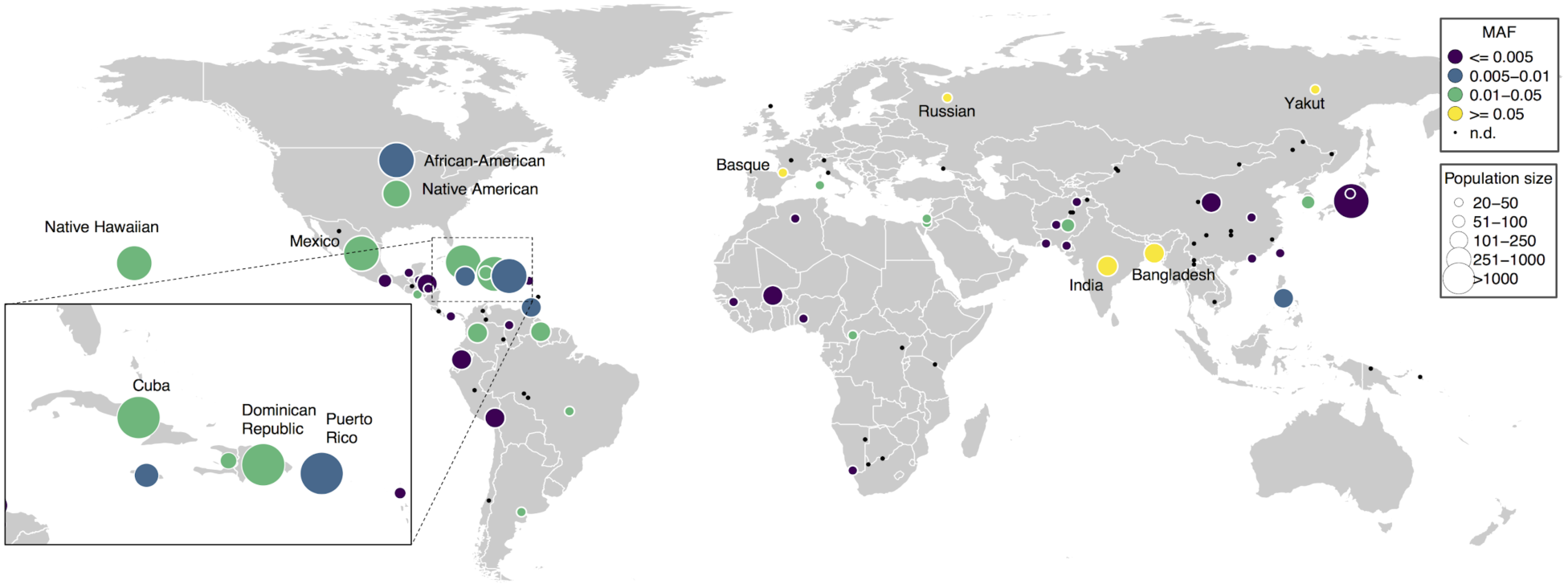
World map of *HCP5*-G frequencies. The histocompatibility protein variant HLA-B*57:01 interacts with the HIV drug abacavir to stimulate a hypersensitivity response. A variant in a gene near HLA-B, HCP5 rs2395029 (G allele), can be used to genotype for the −B*57:01 allele as it is in high linkage disequilibrium (correlation ~0.92 in 1000 Genomes Phase 1). ^45,46,50,51^. This HCP5 tag-SNP segregates within all continental populations of the PAGE study, providing increased resolution of the global haplotype frequency, particularly within Latin America. Above, minor allele (G) frequency is shown. Population size is indicated by the radius of the circle. Black dot (MAF not displayed): population has less than twenty individuals or the variant is a singleton in that population.

## Discussion

Our understanding of the genetics underlying most common traits is derived primarily from individuals of European descent. To address this gap in representation, we assembled the PAGE Study, an initiative to enhance our understanding of the genetics of complex traits through multi-ethnic analyses of a large number of measured phenotypes. We provide study design, methods, and best practices to move beyond the single population bias in GWAS. We perform well-powered, multi-ethnic GWAS to both identify novel signals and better characterize known loci by incorporating and accounting for the systematic role of population structure in genome-phenome architecture. We empirically assess a genomic database of common diseases and traits (GWAS Catalog), and demonstrate that over a quarter of variants in that database show evidence of differential effects across the spectrum of human diversity in PAGE. Furthermore, our results suggest that a majority of GWAS Catalog associations as reported are unlikely to be the causal variant from our fine-mapping in a multi-ethnic population, consistent with differential LD between tagSNPs and functional variants across populations. Finally, we demonstrate that meta-analysis of existing data with a multi-ethnic population (rather than more Europeans) not only reduces the gap in variance explained between European and minority populations, but also significantly improves fine-mapping of known loci.

We have focused on quantitating the scientific value of including diverse populations in the discovery and replication phases of GWAS. Population-specific allele frequencies allow underpowered rare variants in existing GWAS to be more easily detected in different populations, where these variants may have reached a higher frequency facilitating their discovery. 7 Additionally, differential patterns of linkage disequilibrium between populations facilitate fine-mapping within known loci. In both of these contexts, we show that meta-analysis of existing data with a diverse population (~50,000 PAGE individuals) provides added value (novel loci, or better refinement of known loci) over meta-analysis with a similar-sized Europeans-only study, even when combining PAGE with much larger studies. Further, polygenic risk prediction models are an active area of development in human genetics and researchers need to be aware of the limitations of primary discoveries that have not been replicated in multiple populations. ^12^ As we move toward incorporating GWAS-based risk models in clinical care^52^, this, and other recent work^53^, demonstrates that we risk exacerbating health disparities unless diverse, multi-ethnic studies are included. This study also provides evidence that a significant number of novel loci (as well as independent, secondary alleles in known loci) relevant to non-European ancestries remain to be identified, many of which cannot be discovered in European-only study populations due to their low allele frequencies or absence in Europeans. Cumulatively, these results expose several shortcomings that arise from an overreliance on discovery genomics focused on a small number of populations.

As next-generation sequencing, precision medicine, and direct-to-consumer genetic testing become more common, it is critical that the genetics community takes a forward-thinking approach towards the opportunities presented by including diverse populations. The increasing ability to identify rare variants further highlights the necessity to study genetically diverse populations, as rare variation is more likely to be ancestry specific.^4^ And, as we have demonstrated by adding rare, clinically relevant variants to MEGA, these alleles can be highly differentiated across populations. In the United States, the All of Us Research Program (AoURP) embraces the reality that the success of precision medicine requires precision genomics, and therefore emphasizes the recruitment and active participation of underrepresented populations. ^54^ It is in the best interest of our research community to follow suit and take steps to become more inclusive. As world populations become increasingly complex ^55,56^, geneticists and clinicians will be required to evaluate genetic predictors of complex traits in more and more diverse populations. Our current genomic databases are under representative of populations with the greatest health burden or that will ultimately benefit most from this work. This realization, combined with the increased availability of resources for studying diverse populations, means that researchers and funders can no longer afford to ignore non-European populations. Our study provides valuable resources in the design of MEGA and through the sharing of population-specific allele frequencies and analysis approaches, which will provide the motivation to make research in diverse populations a priority in the field of genetics.

## Methods

### Studies

The PAGE Study includes eligible minority participants from four studies. The Women’s Health Initiative (WHI) is a long-term, prospective, multi-center cohort study investigating post-menopausal women’s health in the US, which recruited women from 1993-1998 at 40 centers across the US. WHI participants reporting European descent were excluded from this analysis. The Hispanic Community Health Study / Study of Latinos (HCHS/SOL) is a multi-center study of Hispanic/Latinos with the goal of determining the role of acculturation in the prevalence and development of diseases relevant to Hispanic/Latino health. Starting in 2006, household sampling was used to recruit self-identified Hispanic/Latinos from four sites in San Diego, CA, Chicago, IL, Bronx, NY, and Miami, FL. All SOL Hispanic/Latinos were eligible for this study. The Multiethnic Cohort (MEC) is a population-based prospective cohort study recruiting men and women from Hawaii and California, beginning in 1993, and examines lifestyle risk factors and genetic susceptibility to cancer. Only the African American, Japanese American, and Native Hawaiian participants for MEC were included in this study. The BioMeTM BioBank is managed by the Charles Bronfman Institute for Personalized Medicine at Mount Sinai Medical Center (MSMC). Recruitment began in 2007 and continues at 30 clinical care sites throughout New York City. BioMe participants were African American (25%), Hispanic/Latino, primarily of Caribbean origin (36%), Caucasian (30%), and Others who did not identify with any of the available options (9%). Biobank participants who self-identified as Caucasian were excluded from this analysis. The Global Reference Panel (GRP) was created from Stanford-contributed samples to serve as a population reference dataset for global populations. GRP individuals do not have phenotype data and were only used to aid in the evaluation of genetic ancestry in the PAGE samples. Additional information about each participating study can be found in the Supplementary Information.

### Phenotypes

The 26 phenotypes included in this study were previously harmonized across the PAGE studies. They include: White Blood Cell (WBC) count, C-Reactive Protein (CRP), Mean Corpuscular Hemoglobin Concentration (MCHC), Platelet Count (PLT), High Density Lipoprotein (HDL), Low-Density Lipoprotein (LDL), Total Cholesterol (TC), Triglycerides (TG), glycated hemoglobin (HbA1c), Fasting Insulin (FI), Fasting Glucose (FG), Type II Diabetes (T2D), Cigarettes per Day (CPD), Coffee Consumption, QT interval, QRS interval, PR interval, Systolic Blood Pressure (SBP), Diastolic Blood Pressure (DBP), Hypertension (HT), Body Mass Index (BMI), Waist-to-hip ratio (WHR), Height (HT), Chronic Kidney Disease (CKD), End-Stage Renal Disease (ESRD), and Estimated glomerular filtration rate (eGFR) by the CKD-Epi equation. Single variant association testing was completed for all phenotypes using phenotype-specific models, adjusting by indicators for study, self-identified race/ethnicity as a proxy for cultural background, phenotype-specific standard covariates, and the first 10 PCs. Additional information about phenotype-specific cleaning, exclusion criteria, and the model covariates are included in the Supplementary Information.

### Genotyping

A total of 53,338 PAGE and GRP samples were genotyped on the MEGA array at the Johns Hopkins Center for Inherited Disease Research (CIDR), with 52,878 samples successfully passing CIDR’s QC process. Genotyping data that passed initial quality control at CIDR were released to the Quality Assurance / Quality Control (QA/QC) analysis team at the University of Washington Genetic Analysis Center (UWGAC). The UWGAC further cleaned the data according to previously described methods ^44^, and returned genotypes for 51,520 subjects. A total of 1,705,969 SNPs were genotyped on the MEGA. Quality Control of genotyped variants was completed by filtering through various criteria, including the exclusion of (1) CIDR technical filters, (2) variants with missing call rate >= 2%, (3) variants with more than 6 discordant calls in 988 study duplicates, (4) variants with greater than 1 Mendelian errors in 282 trios and 1439 duos, (5) variants with a Hardy-Weinberg p-value less than 1×10^−4^, (6) SNPs with sex difference in allele frequency >= 0.2 for autosomes/XY, (7) SNPs with sex difference in heterozygosity > 0.3 for autosomes/XY, (8) positional duplicates. Sites were further restricted to chromosomes 1-22, X, or XY, and only variants with available strand information. After SNP QC, a total of 1,402,653 MEGA variants remained for further analyses. (For more details see **Supplementary Information Section 3**)

### Imputation

In order to increase coverage, and thus improve power for fine-mapping loci, all PAGE individuals who were successfully genotyped on MEGA were subsequently imputed into the 1000 Genomes Phase 3 data release^57^. Imputation was conducted at the University of Washington Genetic Analysis Center (GAC). Genotype data which passed the above quality control filters was phased with SHAPEIT2 ^58^ and imputed to 1000 Genomes Phase 3 reference data using IMPUTE version 2.3.2 ^59^. Segments of the genome which were known to harbor gross chromosomal anomalies were filtered out of the final genotype probabilities files. Imputed sites were excluded if the IMPUTE info score was less than 0.4. A total of 39,723,562 imputed SNPs passed quality control measures. (For more details see **Supplementary Information Section 3**)

### Principal Component Analysis

The SNPRelate (Zheng et al. 2012) package in R was used for principal components analysis. (See Supplement for further details.) The relevant principal components (PCs) were selected using scatter plots. Scatter plots, with various PCs on the x- and y-axes, helped to assess the spread of genetic ancestry within with self-identified racial/ethnic clusters. A parallel coordinate plots for the first 10 PCs was generated, where each PAGE individual is represented by a set of line segments connecting his or her PC values. The amount of variance explained diminished with each subsequent PC, and we estimated that the top 10 PCs provided sufficient information to explain the majority of genetic variation in the PAGE study population.

### Genome-Wide Association Testing

All imputed autosomal variants with IMPUTE info score >0.4 (n=39,723,562) were eligible for association testing in phenotype-specific models. An effective sample size (effN) was calculated for each SNP in a given phenotype-specific model, where effN = 2*MAF*(1- MAF)*N*info, where MAF is the minor allele frequency among the set of individuals included in a phenotype-specific model, N is the total sample size for a given phenotype, and info is the SNP’s IMPUTE info score. Variants with an effN less than 30 (continuous phenotypes) or 50 (binary phenotypes), were excluded from the final set of phenotype-specific results. The number of variants analyzed per traits ranged from 21,894,105 to 34,656,550 for continuous phenotypes and 11,665,604 to 28,263,875 for binary phenotypes (**Supplementary Table 1**). QQ plots and lambdas were used to assess genomic inflation in all phenotypes, for which lambdas ranged from 0.98 to 1.15. Single-variant association testing for each phenotype used an additive model that was adjusted by indicators for study, self-identified race/ethnicity, the first 10 PCs, and phenotype-specific covariates. Additional information about the phenotype-specific model covariates and transformations are included in the Supplementary Information. Association testing was completed in both SUGEN and GENESIS programs.

The GENESIS ^19,20^ program is a Bioconductor package made available in R that was developed for large-scale genetic analyses in samples with complex structure including relatedness, population structure, and ancestry admixture. The current version of GENESIS implements both linear and logistic mixed model regression for genome-wide association testing. The software can accommodate continuous and binary phenotypes. The GENESIS package includes the program PC-Relate, which uses a principal component analysis based method to infer genetic relatedness in samples with unspecified and unknown population structure. By using individual-specific allele frequencies estimated from the sample with principal component eigenvectors, it provides robust estimates of kinship coefficients and identity-by-descent (IBD) sharing probabilities in samples with population structure, admixture, and HWE departures. It does not require additional reference population panels or prior specification of the number of ancestral subpopulations.

The SUGEN program ^22^ is a command-line software program developed for genetic association analysis under complex survey sampling and relatedness patterns. It implements the generalized estimating equation (GEE) method, which does not require modeling the correlation structures of complex pedigrees. It adopts a modified version of the “sandwich” variance estimator, which is accurate for low-frequency SNPs. Association testing in SUGEN requires the formation of “extended” families by connecting the households who share first degree relatives or either first- or second-degree relatives. Trait values are assumed to be correlated within families but independent between families. In our experience in analyzing this dataset, it is sufficient to account for first-degree relatedness. The current version of SUGEN can accommodate continuous, binary, and age-at-onset traits. A comparison of p-values produced by SUGEN and GENESIS for all previously identified known loci are included in **Supplementary Figure 11**.

### Conditional Analyses

Phenotype-specific lists of previously identified “known loci” were hand-curated for each phenotype and included SNPs indexed in the GWAS Catalog or identified through non-GWAS high-throughput methods (e.g. Metabochip, Exomechip, Immunochip, etc.). The full known loci lists for each phenotype are available in the Supplementary Table 5. Conditional analyses were conducted for all phenotypes by conditioning on all previously identified known loci on a given chromosome. P-values estimated in conditional analyses are denoted by “P_cond_” in the main text, with the SUGEN conditional results for all novel and secondary findings in **Supplementary Table 3**.

### SNPxPC Effect Heterogeneity by Genetic Ancestry and Self-Identified Race/Ethnicity

We used two approaches to assess effect heterogeneity within PAGE participants. First, we used interaction analyses with models that included variant by PC (SNPxPC) interaction terms for all 10 PCs. The fit of nested models was compared using the F-statistic, where the associated interaction p-value indicated whether the inclusion of the 10 SNPxPC interaction terms improved the model fit compared to a model that lacked the interaction terms. The overall SNPxPC interaction p-values evaluated whether the additional variance explained by variant x genetic ancestry interactions was statistically significant, and represent effect modification driven by genetic ancestry. Interaction p-values for all novel and secondary findings are included in Supplementary Table 3.

For comparison against more traditional (stratified) analysis strategies, all analyses were also run stratified by self-identified race/ethnicity. A minor allele count of at least 5 was required for a stratified model to be run within an ethnic group. The stratified analyses were then meta-analyzed using a fixed-effect model implemented in METAL^49^. I^2^ and chi^2^ heterogeneity p-values were estimated for all meta-analyzed results, and represent effect size heterogeneity driven by self-identified race/ethnicity. The race/ethnicity-specific results, I^2^, and chi^2^ heterogeneity p-values for all novel and secondary findings are included in **Supplementary Table 3**.

### Standardized Effect Size (z’) Analysis

The standardized effect size (z’) analysis for **Figure 2A** was performed as follows. To avoid double-counting of SNPs/loci, we constrained analysis for each trait to (a) the single previous report that (b) did not combine genome-wide genotypes with focused platforms like the metabochip, (c) reported the direction of effect with the allele in the GWAS catalog, and (d) included the maximum total number of individuals after applying criteria (a) and (b). (a) We selected a single manuscript because many traits already have serial meta-analyses published, where earlier publications represent a subset of individuals reported in later publications, so reported effect sizes in the GWAS catalog are not necessarily independent. (b) We excluded meta-analyses using mixtures of agnostic GWAS data (consistent map density across the genome) with focused platforms (e.g. metabochip, oncochip, or exome chip) because the actual sample size varies dramatically across the genome, with overlapping agnostic/focused regions having substantially greater numbers of individuals in the analysis. Sadly, most of these reports fail to specify the sample-size on a per-SNP basis, making it impossible to confidently calculate z’. (c) Starting from the 22 quantitative traits, we found reference studies that explicitly reported the allele associated <u>with direction of effect</u> for 18. Furthermore, in order to be confident that the direction of effect was consistent between PAGE and prior reports, we restricted analysis to asymmetric SNPs (A/C, A/G, C/T and G/T). These criteria yielded 615 previously reported genome-wide significant variants, distributed across the 18 traits (**Supplementary Table 7**). Only 115 of these variants were traditionally genome-wide significant (p<5e-8) and therefore overlap with the SNPxPC heterogeneity analysis.

### Assessing Single-Variant Results

SUGEN association results were used for the identification of novel and secondary findings for all phenotypes. The variant with the smallest p-value in a 1Mb region was considered the “lead SNP”. A lead SNP was considered to be a novel loci if it met the following criteria: 1) the lead SNP was located greater than +/- 500 Kb away from a previously known loci (per the phenotype-specific known loci list); 2) had a SUGEN p-value less than 5×10^−8^; 3) had a SUGEN conditional p-value less than 5×10^−8^ after adjustment for all previously known loci on the same chromosome; and 4) had 2 or more neighboring SNPs (within +/- 500 Kb) with a p-value less than 1×10^−5^. A lead SNP was considered to be a secondary signal in a previously known loci if it met the following criteria: 1) the lead SNP was located within +/- 500 Kb of a previously known loci; 2) had a SUGEN p-value less than 5×10^−8^; and 3) had a SUGEN conditional p-value less than 5×10^−8^ after adjustment for all previously known loci on the same chromosome. Full results for all novel and secondary findings are included in **Supplementary Table 2-3.**

### Effect Size Heterogeneity in the GWAS Catalog

The full GWAS Catalog database was downloaded on December 31, 2016. ^30^ The data were filtered to identify results relevant to any of the 26 PAGE phenotypes, producing a subset of 3,322 unique tagSNPs that were genome-wide significant (p<5×10^−8^) in the GWAS Catalog. The PAGE results for each of the 3,322 GWAS Catalog tagSNPs was examined to first identify the subset of tagSNPs that replicated (p<5×10^−8^) in PAGE unconditioned models. Pairs of replicated tagSNPs within 500,000 base pairs of each other were then merged into loci, in order to count “unique” associated loci. Of the GWAS Catalog tagSNPs that were replicated in PAGE, SNPs that had a Bonferroni corrected SNPxPC interaction heterogeneity p-value (p < 8.71×10^−5^, 0.05/574) were considered to have evidence of effect size heterogeneity between ancestries. Effect heterogeneity was also assessed using PAGE’s multi-ethnic study population by first identifying the “lead SNP” in each locus with the smallest p-value in PAGE, totaling 333 SNPs (302 known loci from the GWAS catalog, plus 31 novel loci discovered in the present analysis). Among the 333 lead SNPs, 24 (7.2%) had a significant Bonferroni corrected SNPxPC interaction heterogeneity p-value (P<1.5×10^−4^, 0.05/333).

### Meta-analysis and Finemapping with GIANT, UKB50k

#### Meta-analysis

We meta-analyzed results for BMI and height in our PAGE multiethnic sample (~50,000 individuals) with the published data from GIANT consortium32,33 which included approximately >250,000 individuals of European descent for each trait. We a comparison for value added, we also conducted a meta-analysis with 50,000 randomly sampled “White-British’’ individuals from the UK Biobank (UKB50k). We conducted GWAS for both PAGE and UKB50k with analogous models for BMI and height traits. Within PAGE and UKB50k, we used the inverse normally transformed residuals for each trait by sex and race/ethnicity, and adjusted for population substructure, age, center, and racial/ethnic groups (if applicable). These methods were similar to what was performed by GIANT, using inverse normal adjusted residuals for each trait outcome. We then meta-analyzed results using a fixed effects model from each PAGE or UKB50k with GIANT separately with METAL software ^60^. We retained only variants available in the combined meta-analyses (for GIANT+PAGE or GIANT+UKB50k) which was approximately 2.5 million. Significance was defined at P-value <5×10-8. Novelty of a locus was defined as +/-500kb from anything known for the respective trait based on the GIANT published results in Locke et al. 2015 (^32,33^. We also required the at least 2 SNPs within a 1Mb results had a P-value <1×10^−5^ to be retained as a significant known or novel locus.

#### Finemapping

We used FINEMAP ^61^ for all finemapping analyses. For each previously-reported locus for height ^33^ and BMI ^32^ in GIANT, a one megabase region was subset, using the summary statistics from GIANT, the PAGE+GIANT meta-analysis, and the UKB50k+GIANT meta-analysis. The linkage disequilibrium for the finemapping was calculated using each individual ancestry from the PAGE sample and using the individuals of European descent from the ARIC study. For weighted linkage disequilibrium that included all ancestries, we weighted each ancestry in PAGE by the actual sample size and added in the ARIC sample but used the sample size from the GIANT consortium by trait. All analyses were run assuming one causal variant. The cumulative 95% credible set was calculated from the estimated posterior probabilities.

### Proportion of variance explained (PVE)

Each PVE analysis considered a single combination of (a) trait, (b) the analysis from which p-values were derived (GIANT, GIANT+PAGE, or GIANT_UKB50k), and (c) the target population in which PVE was calculated (either PAGE or UKB50k). To avoid over-weighting any single region due to LD between multiple associated SNPs, we first defined a “locus” as a contiguous series of genome-wide significant tagSNPs with genome wide significance, where each tagSNP was less than 500k from the next. Then we selected the single SNP within each locus with the smallest p-value in the given analysis (the “best” tagSNP) and calculated PVE for that SNP in the target population. The meta-analysis was effectively limited to allele frequencies greater than 5%, so we used the standard P<5×10^−8^ threshold for significance to define loci.

PVE was calculated for a given SNP using equation 4 from Supplement 1 in Shim et al. ^34^

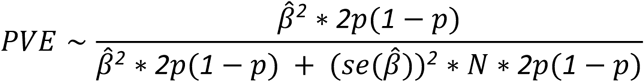

This requires only the estimated effect size (*β̂*) the standard error of the estimate (se(*β̂*)), the allele frequency (p), and the number of samples (N). PVE was then summed across all of the “best” tagSNPs in a given analysis.

### Population allele frequencies of *HCP* rs2395029[G]

These 99 labels were compiled from self-identified ancestry information from the PAGE sample manifest, as well as self-reported country of origin from the Mount Sinai Bio*Me* biobank. Per-population allele frequencies for rs2395029[G] were calculated in PLINK v.1.90 (www.cog-genomics.org/plink/1.9/) ^62^, and results were visualized in R (available at https://github.com/epsorokin/clinical_genetics).

## Acknowledgements

The Population Architecture Using Genomics and Epidemiology (PAGE) program is funded by the National Human Genome Research Institute (NHGRI) with co-funding from the National Institute on Minority Health and Health Disparities (NIMHD). The contents of this paper are solely the responsibility of the authors and do not necessarily represent the official views of the NIH. The PAGE consortium thanks the staff and participants of all PAGE studies for their important contributions. We thank Rasheeda Williams and Margaret Ginoza for providing assistance with program coordination. The complete list of PAGE members can be found at http://www.pagestudy.org.

Assistance with data management, data integration, data dissemination, genotype imputation, ancestry deconvolution, population genetics, analysis pipelines, and general study coordination was provided by the PAGE Coordinating Center (NIH U01HG007419). Genotyping services were provided by the Center for Inherited Disease Research (CIDR). CIDR is fully funded through a federal contract from the National Institutes of Health to The Johns Hopkins University, contract number HHSN268201200008I. Genotype data quality control and quality assurance services were provided by the Genetic Analysis Center in the Biostatistics Department of the University of Washington, through support provided by the CIDR contract. The data and materials included in this report result from collaboration between the following studies and organizations:

## BioMe Biobank

Samples and data of The Charles Bronfman Institute for Personalized Medicine (IPM) BioMe Biobank used in this study were provided by The Charles Bronfman Institute for Personalized Medicine at the Icahn School of Medicine at Mount Sinai (New York). Phenotype data collection was supported by The Andrea and Charles Bronfman Philanthropies. Funding support for the Population Architecture Using Genomics and Epidemiology (PAGE) IPM BioMe Biobank study was provided through the National Human Genome Research Institute (NIH U01HG007417).

## HCHS/SOL

Primary funding support to Dr. North and colleagues is provided by U01HG007416. Additional support was provided via R01DK101855 and 15GRNT25880008. The Hispanic Community Health Study/Study of Latinos was carried out as a collaborative study supported by contracts from the National Heart, Lung, and Blood Institute (NHLBI) to the University of North Carolina (N01-HC65233), University of Miami (N01-HC65234), Albert Einstein College of Medicine (N01-HC65235), Northwestern University (N01-HC65236), and San Diego State University (N01- HC65237). The following Institutes/Centers/Offices contribute to the HCHS/SOL through a transfer of funds to the NHLBI: National Institute on Minority Health and Health Disparities, National Institute on Deafness and Other Communication Disorders, National Institute of Dental and Craniofacial Research, National Institute of Diabetes and Digestive and Kidney Diseases, National Institute of Neurological Disorders and Stroke, NIH Institution-Office of Dietary Supplements.

## MEC

The Multiethnic Cohort study (MEC) characterization of epidemiological architecture is funded through the NHGRI Population Architecture Using Genomics and Epidemiology (PAGE) program (NIH U01 HG007397). The MEC study is funded through the National Cancer Institute U01 CA164973.

## PAGE Global Reference Panel

The Stanford Global Reference Panel was created by Stanford-contributed samples and comprises multiple datasets from multiple researchers across the world designed to provide a resource for any researchers interested in diverse population data on the Multi-Ethnic Global Array (MEGA), funded by the NHGRI PAGE program (NIH U01HG007419). The authors thank the researchers and research participants who made this dataset available to the community. The specific datasets are:

Mexico: Samples of indigenous origin in Oaxaca were kindly provided by co-authors, and Samuel Canizales Quinteros and Victor Acuña Alonzo. Peru: Individuals from a primarily Quechuan and Aymaran-speaking community in Puno were kindly provided with funding support from the Burroughs Welcome Fund. Rapa Nui (Easter Island, Chile): Samples were kindly provided with funding from the Charles Rosenkranz Prize for Health Care Research in Developing Countries and the International Center for Genetic Engineering and Biotechnology (ICGEB) Grant CRP/MEX15- 04_EC awarded to A.M.-E. South Africa: Samples of KhoeSan individuals from the ‡Khomani and Nama communities were kindly provided with funding from the Morrison Institute for Population and Resource Studies. Honduras and Colombia: Samples from communities in Honduras and Colombia were kindly provided by co-authors, Edwin Herraro-Paz (Universidad Católica de Honduras, San Pedro Sula, Honduras), Alvaro Mayorga (Universidad Católica de Honduras, San Pedro Sula, Honduras), Luis Caraballo (University of Cartagena), Javier Marrugo (university of Cartagena) Additional global samples: The following datasets are open access and available through the lab website of Carlos Bustamante (https://bustamantelab.stanford.edu/). The Human Genome Diversity Panel (HGDP-CEPH) is a group of cell lines maintained by the Centre d’Étude du Polymorphisme Humain, Fondation Jean Dausset (Paris, France) comprising 52 diverse populations across the world (Africa, Near East, Europe, South Asia, Central Asia, East Asia, Oceania and the Americas). Additional information on these datasets can be found on the CEPH website (http://www.cephb.fr/en/hgdp_panel.php), or originally at http://www.ncbi.nlm.nih.gov/pubmed/11954565 and http://www.ncbi.nlm.nih.gov/pubmed/12493913, with numerous subsequent publications. Samples were filtered to include the H952 unrelated individuals as published here: http://www.ncbi.nlm.nih.gov/pubmed/17044859. Also available on the Bustamante Lab website is genotype data for the Maasai from Kinyawa, Kenya (MKK) samples maintained by the Coriell Institute for Medical Research (https://catalog.coriell.org/1/NHGRI/Collections/HapMap-Collections/Maasai-in-Kinyawa-Kenya-MKK) and genotyped as part of the International HapMap Project Phase 3(http://hapmap.ncbi.nlm.nih.gov/, http://www.sanger.ac.uk/resources/downloads/human/hapmap3.html). We have genotyped a subset of unrelated individuals using the filters recommended in http://www.ncbi.nlm.nih.gov/pubmed/20869033.

## WHI

Funding support for the “Exonic variants and their relation to complex traits in minorities of the WHI” study is provided through the NHGRI PAGE program (NIH U01HG007376). The WHI program is funded by the National Heart, Lung, and Blood Institute, National Institutes of Health, U.S. Department of Health and Human Services through contracts HHSN268201100046C, HHSN268201100001C, HHSN268201100002C, HHSN268201100003C, HHSN268201100004C, and HHSN271201100004C. The authors thank the WHI investigators and staff for their dedication, and the study participants for making the program possible. A listing of WHI investigators can be found at: https://www.whi.org/researchers/Documents%20%20Write%20a%20Paper/WHI%20Investigator%20Short%20List.pdf

## Individual Acknowledgements

KKN was supported by the Cancer Prevention Training Grant in Nutrition, Exercise and Genetics R25CA094880 from the National Cancer Institute. CRG was supported by NHGRI training grant T32 HG000044. HMH was supported by NHLBI training grant T32 HL007055. AEJ was supported by NIH 5K99HL130580-02 and NIH L60 MD008384-02. KLY supported by NCATS KL2TR001109. JMK was supported by KL2TR000421. RWW was supported by NIH 5T32HD049311-07. D-YL was supported by R01CA082659, R01GM047845, and P01CA142538. LFR was supported by NICHD training grant T32 HD007168 and P2C HD050924. TAT was supported by P01GM099568.

## Author Contributions

Overall project supervision and management: ED, J-LA, LRW, RSJ, LAH, SB, CH, CK, LLM, RJFL, TM, KEN, UP, EEK, CSC. Genotyping and quality control: GLW, JH, CRG, NZ, SB, JMK, EPS, KV, GMB, RWW, CS, MHP, MF, CDB, LCP, JR, KD, MPC, XS, CAL, CCL, RD, GN, EB, SCN, CK, UP, EEK, CSC. Phenotype harmonization: MG, KKN, JH, HMH, YMP, AEJ, CJH, CLW, CLA, KLY, MAR, NZ, SB, JMK, IC, VWS, GMB, CS, AV, MHP, GH, LFR, MF, APR, LRW, YL, S-SLP, CPC, RD, GN, EB, SB, CK, LLM, UP, EEK. Association analyses: GLW, MG, KKN, RT, JH, CRG, HMH, YMP, AEJ, BML, CJH, CLW, CLA, KLY, MAR, SB, JMK, IC, VWS, EPS, GMB, MV, YL, D-YL. TAT, J-LA, DOS, YL, S-SLP, CK, UP, EEK, CSC. Manuscript preparation: GLW, MG, KKN, RT, JH, CRG, HMH, YMP, AEJ, BML, CJH, CLW, CLA, KLY, MAR, JMK, IC, VWS, EPS, RWW, AV, LH, D-YL, GH, APR, TAT, DOS, RSJ, LAH, RD, GN, EAS, SB, CH, CK, LLM, RJFL, TM, KEN, UP, EEK, CSC.

## Data Availability

Individual-level phenotype and genotype data are available through dbGaP at https://www.ncbi.nlm.nih.gov/projects/gap/cgi-bin/study.cgi?study_id=phs000356. Allele frequency data will be available for all genotyped sites on dbSNP (https://www.ncbi.nlm.nih.gov/projects/SNP/) and the University of Chicago Geography of Genetic Variants Browser (http://popgen.uchicago.edu/ggv/). Clinically-relevant variant frequency data will also be available through ClinGen.

## Bibliography

1. Need, A. C. & Goldstein, D. B. Next generation disparities in human genomics: concerns and remedies. Trends Genet. 25, 489–494 (2009).

2. Bustamante, C. D., Burchard, E. G. & De la Vega, F. M. Genomics for the world. Nature 475, 163–165 (2011).

3. Popejoy, A. B. & Fullerton, S. M. Genomics is failing on diversity. Nature 538, 161–164 (2016).

4. Gravel, S. et al. Demographic history and rare allele sharing among human populations. Proc Natl Acad Sci USA 108, 11983–11988 (2011).

5. SIGMA Type 2 Diabetes Consortium et al. Association of a low-frequency variant in HNF1A with type 2 diabetes in a Latino population. JAMA 311, 2305–2314 (2014).

6. Gudmundsson, J. et al. A study based on whole-genome sequencing yields a rare variant at 8q24 associated with prostate cancer. Nat. Genet. 44, 1326–1329 (2012).

7. Moltke, I. et al. A common Greenlandic TBC1D4 variant confers muscle insulin resistance and type 2 diabetes. Nature 512, 190–193 (2014).

8. Kenny, E. E. et al. Melanesian blond hair is caused by an amino acid change in TYRP1. Science 336, 554 (2012).

9. Manning, A. et al. A Low-Frequency Inactivating AKT2 Variant Enriched in the Finnish Population Is Associated With Fasting Insulin Levels and Type 2 Diabetes Risk. Diabetes 66, 2019–2032 (2017).

10. Han, Y. et al. Prostate cancer susceptibility in men of african ancestry at 8q24. J Natl Cancer Inst 108, (2016).

11. Carlson, C. S. et al. Generalization and dilution of association results from European GWAS in populations of non-European ancestry: the PAGE study. PLoS Biol. 11, e1001661 (2013).

12. Martin, A. R. et al. Human Demographic History Impacts Genetic Risk Prediction across Diverse Populations. Am. J. Hum. Genet. 100, 635–649 (2017).

13. Liao, Y. et al. Surveillance of health status in minority communities - Racial and Ethnic Approaches to Community Health Across the U.S. (REACH U.S.) Risk Factor Survey, United States, 2009. MMWR Surveill. Summ. 60, 1–44 (2011).

14. Oh, S. S. et al. Diversity in clinical and biomedical research: A promise yet to be fulfilled. PLoS Med. 12, e1001918 (2015).

15. Carlson, C. S. Ethnicity: Diversity is future for genetic analysis. Nature 540, 341 (2016).

16. Matise, T. C. et al. The Next PAGE in understanding complex traits: design for the analysis of Population Architecture Using Genetics and Epidemiology (PAGE) Study. Am. J. Epidemiol. 174, 849–859 (2011).

17. Dastani, Z. et al. Novel loci for adiponectin levels and their influence on type 2 diabetes and metabolic traits: a multi-ethnic meta-analysis of 45,891 individuals. PLoS Genet. 8, e1002607 (2012).

18. Demenais, F. et al. Multiancestry association study identifies new asthma risk loci that colocalize with immune-cell enhancer marks. Nat. Genet. 50, 42–53 (2018).

19. Conomos, M. P., Miller, M. B. & Thornton, T. A. Robust inference of population structure for ancestry prediction and correction of stratification in the presence of relatedness. Genet. Epidemiol. 39, 276–293 (2015).

20. Conomos, M. P., Reiner, A. P., Weir, B. S. & Thornton, T. A. Model-free Estimation of Recent Genetic Relatedness. Am. J. Hum. Genet. 98, 127–148 (2016).

21. Conomos, M. P. et al. Genetic diversity and association studies in US hispanic/latino populations: applications in the hispanic community health study/study of latinos. Am. J. Hum. Genet. 98, 165–184 (2016).

22. Lin, D.-Y. et al. Genetic association analysis under complex survey sampling: the Hispanic Community Health Study/Study of Latinos. Am. J. Hum. Genet. 95, 675–688 (2014).

23. Lin, D. Y. & Zeng, D. On the relative efficiency of using summary statistics versus individual-level data in meta-analysis. Biometrika 97, 321–332 (2010).

24. Fadista, J., Manning, A. K., Florez, J. C. & Groop, L. The (in)famous GWAS P-value threshold revisited and updated for low-frequency variants. Eur. J. Hum. Genet. 24, 1202–1205 (2016).

25. Liu, J. Z. et al. Association analyses identify 38 susceptibility loci for inflammatory bowel disease and highlight shared genetic risk across populations. Nat. Genet. 47, 979–986 (2015).

26. Okada, Y. et al. Genetics of rheumatoid arthritis contributes to biology and drug discovery. Nature 506, 376–381 (2014).

27. Torgerson, D. G. et al. Meta-analysis of genome-wide association studies of asthma in ethnically diverse North American populations. Nat. Genet. 43, 887–892 (2011).

28. DIAbetes Genetics Replication And Meta-analysis (DIAGRAM) Consortium et al. Genome-wide trans-ancestry meta-analysis provides insight into the genetic architecture of type 2 diabetes susceptibility. Nat. Genet. 46, 234–244 (2014).

29. Waters, K. M. et al. Consistent association of type 2 diabetes risk variants found in europeans in diverse racial and ethnic groups. PLoS Genet. 6, e1001078 (2010).

30. MacArthur, J. et al. The new NHGRI-EBI Catalog of published genome-wide association studies (GWAS Catalog). Nucleic Acids Res. 45, D896–D901 (2017).

31. Marigorta, U. M. & Navarro, A. High trans-ethnic replicability of GWAS results implies common causal variants. PLoS Genet. 9, e1003566 (2013).

32. Locke, A. E. et al. Genetic studies of body mass index yield new insights for obesity biology. Nature 518, 197–206 (2015).

33. Wood, A. R. et al. Defining the role of common variation in the genomic and biological architecture of adult human height. Nat. Genet. 46, 1173–1186 (2014).

34. Shim, H. et al. A multivariate genome-wide association analysis of 10 LDL subfractions, and their response to statin treatment, in 1868 Caucasians. PLoS ONE 10, e0120758 (2015).

35. Bien, S. A. et al. Strategies for enriching variant coverage in candidate disease loci on a multiethnic genotyping array. PLoS ONE 11, e0167758 (2016).

36. Lacy, M. E. et al. Association of sickle cell trait with hemoglobin a1c in african americans. JAMA 317, 507–515 (2017).

37. Lin, C.-N. et al. Effects of hemoglobin C, D, E, and S traits on measurements of HbA1c by six methods. Clin. Chim. Acta 413, 819–821 (2012).

38. Mongia, S. K. et al. Effects of hemoglobin C and S traits on the results of 14 commercial glycated hemoglobin assays. Am. J. Clin. Pathol. 130, 136–140 (2008).

39. Roberts, W. L. et al. Effects of hemoglobin C and S traits on glycohemoglobin measurements by eleven methods. Clin. Chem. 51, 776–778 (2005).

40. Mallal, S. et al. HLA-B*5701 screening for hypersensitivity to abacavir. N. Engl. J. Med. 358, 568–579 (2008).

41. Sousa-Pinto, B. et al. Pharmacogenetics of abacavir hypersensitivity: A systematic review and meta-analysis of the association with HLA-B*57:01. J. Allergy Clin. Immunol. 136, 1092–4.e3 (2015).

42. Hetherington, S. et al. Hypersensitivity reactions during therapy with the nucleoside reverse transcriptase inhibitor abacavir. Clin. Ther. 23, 1603–1614 (2001).

43. Illing, P. T. et al. Immune self-reactivity triggered by drug-modified HLA-peptide repertoire. Nature 486, 554–558 (2012).

44. Dean, L. in Medical Genetics Summaries (eds. Pratt, V., McLeod, H., Dean, L., Malheiro, A. & Rubinstein, W.) (National Center for Biotechnology Information (US), 2012).

45. Martin, M. A. et al. Clinical Pharmacogenetics Implementation Consortium Guidelines for HLA-B Genotype and Abacavir Dosing: 2014 update. Clin. Pharmacol. Ther. 95, 499–500 (2014).

46. Pappas, D. J. et al. Significant variation between SNP-based HLA imputations in diverse populations: the last mile is the hardest. Pharmacogenomics J. 18, 367–376 (2018).

47. Henn, B. M. et al. Hunter-gatherer genomic diversity suggests a southern African origin for modern humans. Proc Natl Acad Sci USA 108, 5154–5162 (2011).

48. Baker, J. L., Shriner, D., Bentley, A. R. & Rotimi, C. N. Pharmacogenomic implications of the evolutionary history of infectious diseases in Africa. Pharmacogenomics J. 17, 112–120 (2017).

49. Nemat-Gorgani, N. et al. Different Selected Mechanisms Attenuated the Inhibitory Interaction of KIR2DL1 with C2+ HLA-C in Two Indigenous Human Populations in Southern Africa. J. Immunol. 200, 2640–2655 (2018).

50. Colombo, S. et al. The HCP5 single-nucleotide polymorphism: a simple screening tool for prediction of hypersensitivity reaction to abacavir. J. Infect. Dis. 198, 864–867 (2008).

51. Sanchez-Giron, F. et al. Association of the genetic marker for abacavir hypersensitivity HLA-B*5701 with HCP5 rs2395029 in Mexican Mestizos. Pharmacogenomics 12, 809–814 (2011).

52. Khera, A. V. et al. Genome-wide polygenic scores for common diseases identify individuals with risk equivalent to monogenic mutations. Nat. Genet. 50, 1219–1224 (2018).

53. Lee, J. J. et al. Gene discovery and polygenic prediction from a genome-wide association study of educational attainment in 1.1 million individuals. Nat. Genet. 50, 1112–1121 (2018).

54. Collins, F. S. & Varmus, H. A new initiative on precision medicine. N. Engl. J. Med. 372, 793–795 (2015).

55. Colby, S. L. & Ortman, J. M. Projections of the Size and Composition of the U.S. Population: 2014 to 2060. (United States Census Bureau, 2015).

56. -United Nations Population Fund. State of World Population 2016. (2016). at <http://www.unfpa.org/swop>

57. 1000 Genomes Project Consortium et al. A global reference for human genetic variation. Nature 526, 68–74 (2015).

58. Delaneau, O., Marchini, J. & Zagury, J.-F. A linear complexity phasing method for thousands of genomes. Nat. Methods 9, 179–181 (2011).

59. Howie, B. N., Donnelly, P. & Marchini, J. A flexible and accurate genotype imputation method for the next generation of genome-wide association studies. PLoS Genet. 5, e1000529 (2009).

60. Willer, C. J., Li, Y. & Abecasis, G. R. METAL: fast and efficient meta-analysis of genomewide association scans. Bioinformatics 26, 2190–2191 (2010).

61. Benner, C. et al. FINEMAP: efficient variable selection using summary data from genome-wide association studies. Bioinformatics 32, 1493–1501 (2016).

62. Chang, C. C. et al. Second-generation PLINK: rising to the challenge of larger and richer datasets. Gigascience 4, 7 (2015).

